# Context-dependent spatial sorting of dispersal-related traits in the invasive starlings (*Sturnus vulgaris*) of South Africa and Australia

**DOI:** 10.1101/342451

**Authors:** David J. Phair, Johannes J. Le Roux, Cecile Berthouly-Salazar, Vernon Visser, Bettine Jansen van Vuuren, Adam P.A. Cardilini, Cang Hui

## Abstract

Species undergoing range expansion frequently experience increased dispersal rates, especially among invasive alien species. Such increased dispersal rates have been attributed to ‘spatial sorting’, where traits enhancing dispersal assort towards the expanding range edge while traits enhacing competitiveness are favoured within the core range. To date no single study has compared patterns of spatial sorting across multiple continents for the same species. Here we compared patterns of spatial sorting in *Sturnus vulgaris*, the European starling (hereafter referred to as starlings), in its invasive ranges in South Africa and Australia. Starlings have experienced similar residence times in these two countries. Using multi-scale pattern analyses and generalized additive models, we determine whether dispersal and foraging traits (i.e. the morphological attributes of wings and bills) were sorted along the distance from introduction site. We found apparent patterns of spatial sorting in Australia, but not in South Africa. This difference may be attributed to differences in dispersal rates, clinal variation, environmental heterogeneity, and thus population demography on the two continents. Genetic data suggests that starlings in South Africa have experienced frequent long distance dispersal events, which could have diluted or overridden patterns of spatial sorting.

## Introduction

Many species are experiencing range shifts, largely due to human-mediated factors such as climate change (see Parmesan and Yohe 2003 for review, Peterson 2003, Malcolm et al. 2006, but see Petitpierre et al. 2012) as well as intentional and unintentional species movements resulting in biological invasions (Broennimann et al. 2007; Hui and Richardson 2017). Range shifts resulting from climate change normally manifest with species tracking their climatic niches along latitudinal or altitudinal gradients (Parmesan et al. 1999; Chevin et al. 2010; Kerr et al. 2015). In contrast, invasive species expand their ranges after successful human-mediated introduction and establishment in novel environments, often lacking the abiotic and biotic contraints to range expansion that they experienced in their native ranges (Colautti et al. 2004; Shwartz et al. 2009; Parker et al. 2018). Species invasions therefore represent excellent models for assessing the mechanisms underpinning rapid range expansion within contemporary timescales (Arim et al. 2006) especially under novel environmental conditions.

Increased dispersal rates have been detected for some invasive species (Veit and Lewis 1996; Urban et al. 2008; Hui and Richardson 2017) and dispersal-linked traits are often favoured within such rapidly exapanding populations (Thomas et al. 2001). Explanations for increased spread rates include frequent long distance dispersal (Clark 1998), density dependant dispersal (Kot et al. 1996), temporal dispersal variability (Ellner and Schreiber 2012), and spatial sorting (Shine et al. 2011). Spatial sorting is the process whereby effective dispersal phenotypes accumulate at the leading edge of the invasion, which then results in assortative mating and subsequent enhancement of traits linked to dispersal. Simultaneously low intraspecific competition is expected at the leading edge due to low population densities, resulting in relatively higher individual fitness when compared to individuals from the range core (Brown et al. 2013). Spatial sorting plus elevated fitness at the leading edge have been proposed to work in synergy, termed spatial selection, which can actively select for enhanced dispersal at the leading edge of an invasion or range expansion (Phillips et al. 2008).

Although spatial sorting affects dispersal in novel environments, alternative forces will act alongside or against it once the populations have become established. These include amongst others; sexual selection, local adaptation and competition (Holt et al. 2005; Blackburn et al. 2009; Phillips 2009; Perkins 2012; Perkins et al. 2013). This may lead to boosted investment in traits for resource competition or reproduction in the novel range such as the bill morphology of non-native birds which affects food acquisition (Radford and Du Plessis 2003; Berthouly-Salazar et al. 2012). Consequently, spatial sorting should primarily act on dispersal traits at the expanding range front, whilst selection on foraging and reproductive traits will increasingly result from environmental gradients at the core range. Theoretical studies have identified trade-offs between dispersal and traits linked to competition (Burton et al. 2010) such as aggression (Duckworth 2008), fecundity (Kisdi 2004), lifespan (Dytham and Travis 2006), niche breadth (Kisdi 2002), body condition (Gyllenberg et al. 2008), body size (Kisdi et al. 2012) and self-compatibility in plants (Cheptou and Massol 2009). For instance, despite obvious selection advantages in increased spread rates during invasions, some introduced species suffer from Allee effects (see Taylor and Hastings 2005 for review). In these cases greater spread rates could lead to establishment failure due to difficulties in locating mates in low-density loci at the leading edge (Robinet and Liebhold 2009). The existence of such trade-offs inevitably exacerbates the variation in traits across the full range of a species.

Species invasions represent excellent models for assessing the mechanisms underpinning range expansion within contemporary timescales, as they are usually charaterised by rapid rage expansions (Arim et al. 2006) and occur under novel environmental conditions. Spatial sorting has been detected in various traits linked to dispersal and performance of many invasive species (Phillips 2009; Berthouly-Salazar et al. 2012; Lombaert et al. 2014; Cobben et al. 2015). The paradigmatic invasive cane toad (*Bufo marinus*) in Australia displays longer legs at the range front (Phillips et al. 2008), moves more frequently and in straighter lines (Brown et al. 2014) when compared to individuals at the core of the species’ range, leading to increased dispersal ability at the range front (see Rollins et al. 2015 for review). These observations have been repeatedly reported in other systems, including wing morphology in birds (Simmons and Thomas 2004; Hughes et al. 2007; Baldwin et al. 2010), feet size in mammals (Forsman et al. 2011), dispersal endurance in amphibians (Llewelyn et al. 2010), and seed morphology of plants (Cwynar and MacDonald 1987). Morphological traits that indirectly affect dispersal may also be affected by spatial sorting, such as increased migration speeds (Møller 2010) and brain size relative to body size (Sol et al. 2005; Amiel et al. 2011). Spatial sorting can also act on behavioural traits. For instance, aggression in western bluebirds (*Sialia mexicana*) is linked to more effective dispersal and colonisation ability (Duckworth 2008). Similarly, Rodriguez et al. (2010) showed that the European starling, *Sturnus vulgaris*, can display increased attention to social cues at their range edge compared to individuals from the core. For insects, increased numbers of dispersive females and decreased worker production have been detected at the range edge of the equatorial ant *Petalomyrmex phylax*, suggesting that spatial sorting can affect population structuring (Léotard et al. 2009).

However, to date few analyses have assessed spatial sorting across separate invasions of the same taxon to determine how invasion context may affect spatial sorting. Determining patterns of spatial sorting for multiple and independent invasions of the same species may provide insights into the mechanisms and factors affecting this process. Invasions are highly context dependent and it is therefore expected that environmental and temporal context will impact patterns of spatial sorting. The environmental context includes factors such as environmental heterogeneity and habitat quality (Melbourne et al. 2007), while the temporal context includes factors of introduction history, residence time (Kolar and Lodge 2001; Hayes and Barry 2008), and propagule pressure (see Simberloff 2009 for review), all of which can strongly affect invasion dynamics and success. Even under the same condition, invasion dynamics hardly repeat themselves, creating high unpredictability (Melbourne and Hastings 2009).

Invasive European starlings, *Sturnus vulgaris*, represent a good study system to disentangle the influence of environmnetal vs. temporal variation of patterns of spatial sorting. The species occurs naturally across most of Eurasia and is listed in the top 100 most invasive species (Lowe et al. 2000), with 80% of introduced populations having become established (Sol et al. 2002). It is now established and considered invasive in parts of Australia, New Zealand, North America, Polynesia, South Africa and South America (Long 1981). The three main invaded ranges are in Australia, South Africa, and the USA. In the USA 60 birds were introduced, the rate of range expansion was intially estimated at 11.2 km/y, but has since increased to 51.2km/y (Okubo 1988). Similarly in South Africa, while less than 20 birds were introduced in the late 19th century (Harrison and Cherry 1997), initial spread rates were low (6.1 km/y) and increased over time to 25.7 km/y (Hui et al. 2012). On the other hand, in Australia 89 starlings were introduced to Adelaide in late 1850s and has spread at a constant rate of 20.7 km/y (Hui and Richardson 2017) and over vast distances into western Australia (Woolnough et al. 2005), it has also been suggested that there may be several introduction sites within Australia, with Melbourne as the initial introduction site and possibly being a subsequent source population for introductions at other locations (Long 1981).

Given the similar residence times of starlings in South Africa and Australia and the similarity of climatic conditions between the two regions, starling invasions in these countries pose a unique opportunity to investigate and compare the effects of range expansion on morphological traits. Here, we investigate whether spatial sorting of dispersal-linked morphological traits occurred within South African and Australian starling populations, and whether the same morphological traits responded similarly to range expansion in both countries. We further examine whether foraging traits have also sorted spatially. Finally, we explore the potential environmental drivers of observed spatial structures of dispersal and foraging traits in the two continents. Answers to these questions could help to elucidate both patterns and processes behind boosted range expansion observed in many invasive species.

## Material and Methods

### Data

This study made use of two separate datasets. In South Africa the systematic sampling of live birds by hunters was carried out in May 2011 (see Berthouly-Salazar et al. 2013) resulting in 202 individual birds sampled from 35 localities, roughly covering 270,000 km^2^ (Fig. 1A). The Australian data were collected in 2011 and 2012 using a combination of ‘modified walk-in traps’ and hunters. Australian samples were collected to maximise the latitudinal gradient coverage, as well as being collected on transects along the coast (core) and the range edge. This resulted in 444 individual birds sampled from 25 locations, roughly covering 640,000 km^2^ (Fig. 1B).

**Figure 1:**
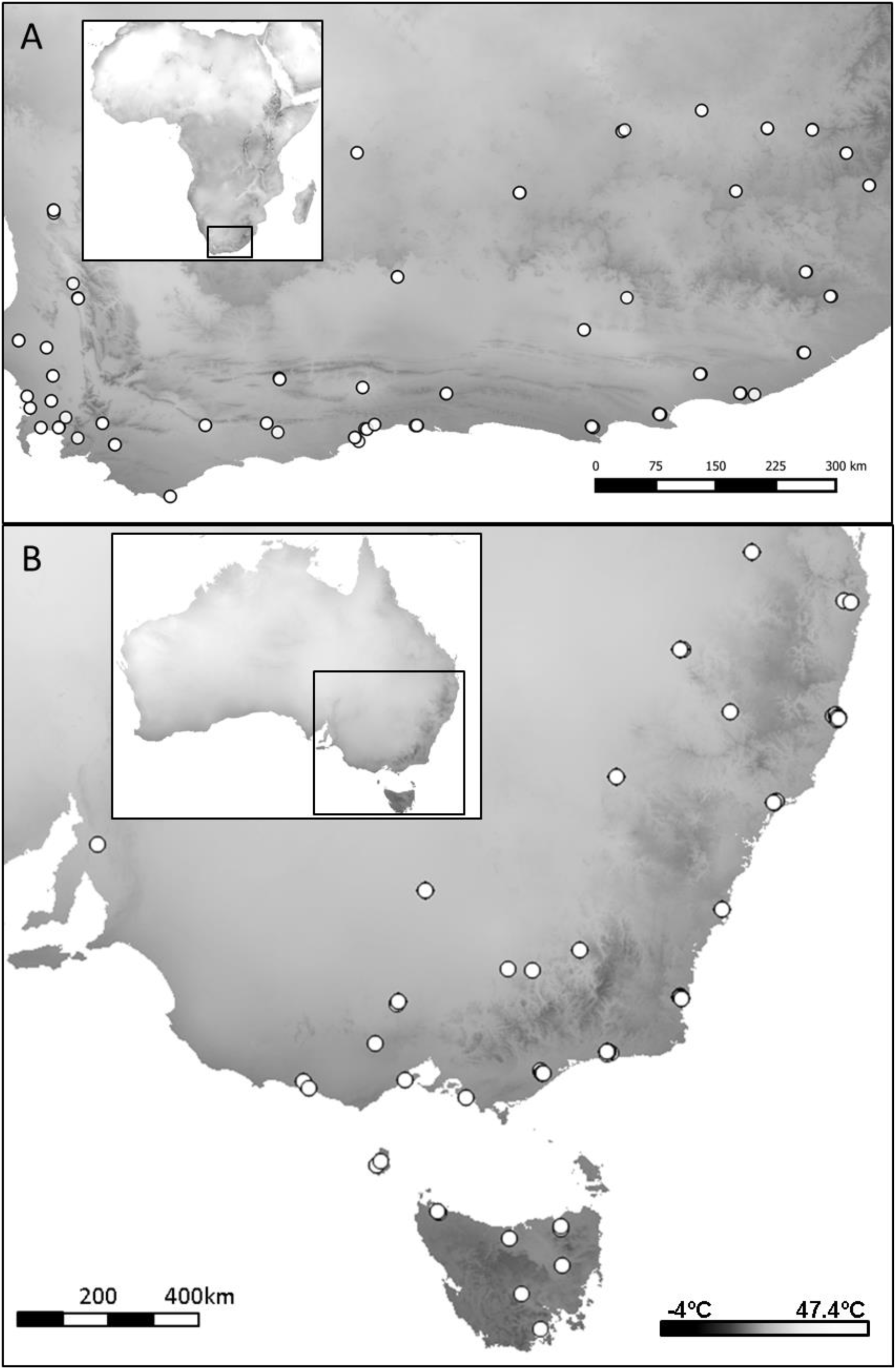
Sampling localities of starlings in South Africa (A) and Australia (B) with an overlay of mean summer maximum temperature (ranging from −4°C to 47.4°C)

All adult birds from both countries were weighed (0.001 g precision) and dissected to determine the sex. A suite of morphological traits (Table S1) were measured and recorded for all birds, including tarsus length, wing length, bill length, bill width, bill depth and head length. We then performed all subsequent analyses using morphological measures corrected for body size. This was accomplished by conducting a Principle Component Analysis (PCA) from which PC1 and PC2 can be seen as size and shape variables respectively; each morphological trait was then regressed against PC1 to calculate size-corrected residues in a generalized linear model. To circumvent statistical dependency, each morphological trait was regressed against the PC1 scores from a PCA of all other traits sans the trait in question and the resultant residuals were employed in all subsequent analyses (Berthouly-Salazar et al. 2012). For both datasets we calculated wing loading via the residual of the regression of *log*_10_[*wing length*^2^] *by log*_10_[*body weight*] (Marchetti et al. 1995) and calculated the following ratios: bill ratio (bill length/bill width), head ratio (head length/weight) and tarsus ratio (tarsus length/ weight). A Kruskal-wallis test was run to determine whether traits differed significantly between adult males, adult females and juveniles for South Africa and Australia (Table 1).

**Table 1:** Measurements (mean ±SD) of morphological traits assessed for adult male, adult female, and juvenile starlings in Asutralia and South Africa.

| Traits | South Africa |  |  | Australia |  |  |
| --- | --- | --- | --- | --- | --- | --- |
|  | Males | Females | Juveniles | Males | Females | Juvenile |
|  | N=66 | N=42 | N=94 | N=66 | N=164 | N=215 |
| Bill Depth[mm] | 7.18 ±<br>0.44<sup>*J</sup> | 6.93±<br>0.35 | 7.14 ±<br>0.39<sup>*M</sup> | 5.82±<br>0.36<sup>*M</sup> | 5.81±<br>0.33<sup>*</sup> | 6.05±<br>0.46<sup>*F</sup> |
| Bill Length[mm] | 21.69 ±<br>1.07<sup>*F</sup> | 20.96 ±<br>1.12<sup>*</sup> | 21.91 ±<br>1.63<sup>*F</sup> | 25.59±<br>1.11<sup>*</sup> | 26.29 ±<br>1.13<sup>*</sup> | 27.18±1<br>.03<sup>*</sup> |
| Bill Width[mm] | 7.11 ±<br>0.43<sup>*J</sup> | 7.07 ±<br>0.47 | 7.41 ±<br>0.46<sup>*M</sup> | 6.20±<br>0.31 | 6.36 ±<br>0.29 | 6.48±0.<br>30 |
| Head Length[mm] | 31.29 ±<br>3.56<sup>*J</sup> | 31.98 ±<br>0.78<sup>*J</sup> | 32.14 ±<br>1.57<sup>*</sup> | 26.48±<br>3.09<sup>*J</sup> | 27.27 ±<br>1.16 | 27.54±0<br>.80<sup>*M</sup> |
| Tarsus Length[mm] | 29.54 ±<br>0.93<sup>*J</sup> | 29.25 ±<br>0.88<sup>*J</sup> | 29.90±<br>0.84<sup>*</sup> | 28.26±<br>0.91 | 28.19 ±<br>0.89 | 28.61±0<br>.83 |
| Wing Length[mm] | 130.55 ±<br>4.61<sup>*J</sup> | 129.00 ±<br>2.52<sup>*J</sup> | 130.68 ±<br>10.74<sup>*</sup> | 125.10±<br>3.81<sup>*</sup> | 129.09 ±<br>2.97<sup>*</sup> | 132.18±<br>2.83<sup>*</sup> |
| Weight[g] | 77.46 ±<br>5.92<sup>*J</sup> | 79.94 ±<br>6.27<sup>*J</sup> | 83.38±5.3<br>1<sup>*</sup> | 73.95 ±<br>6.62<sup>*J</sup> | 74.98 ±<br>5.58<sup>*J</sup> | 79.97±4<br>.78<sup>*</sup> |
| BR (bill length to<br>width Ratio) | 3.06±<br>0.22<sup>*</sup> | 2.98 ±<br>0.29<sup>*M</sup> | 2.97±<br>0.26<sup>*M</sup> | 4.13 ±<br>0.16 | 4.14 ±<br>0.26<sup>*J</sup> | 4.20±0.<br>25<sup>*F</sup> |
| HR (Head to body<br>Ratio) | 0.41 ±<br>0.055<sup>*J</sup> | 0.40 ±<br>0.027<sup>*J</sup> | 0.39±<br>0.030<sup>*</sup> | 0.36 ±<br>0.052<sup>*J</sup> | 0.37±<br>0.028<sup>*J</sup> | 0.35±0.<br>021<sup>*</sup> |
| TR (Tarsus to body<br>Ratio) | 0.38 ±<br>0.030<sup>*</sup> | 0.37 ±<br>0.027<sup>*M</sup> | 0.36±<br>0.021<sup>*M</sup> | 0.38 ±<br>0.032<sup>*J</sup> | 0.38±<br>0.026<sup>*J</sup> | 0.36±0.<br>020<sup>*</sup> |
| Wing Loadings<br>(Residual) | 0.0094 ±<br>0.027<sup>*F</sup> | -0.0047 ±<br>0.019<sup>*</sup> | -0.0054±<br>0.13<sup>*F</sup> | -0.027 ±<br>0.025<sup>*</sup> | -0.0019 ±<br>0.022<sup>*</sup> | 0.010±0<br>.019<sup>*</sup> |

\*indicate significant deviation from the other cases within region unless specified, in which case the differ only from the group specified

The environmental variables that were used included climatic variables attained from the WorldClim database (Hijmans et al. 2005): mean winter precipitation, mean summer precipitation, mean summer maximum temperature and mean winter minimum temperature. For habitat variables we made use of altitude, Normalised Difference Vegetation Index (NDVI) (Lobo 2007), and the distance from the initial introduction site (South Africa: Cape Town; Australia: Melbourne) to each sampled population. Human impact was obtained from the Last of the Wild project’s “Global Human Influence Index” (Sanderson et al. 2002), which is a value bounded between 0 and 72 ranging from zero human influence to complete human influence across nine datasets (Table S2).

### Statistical Analysis

Due to possible researcher bias, we performed all data analyses on the South African and Australian datasets separately. As an initial assesment of differences between core and range populations a Mann-Whitney U test was conducted on the 50 starlings closest to introduction site (representative of the core) and the 50 starlings furthest from introduction site (representative of the edge) in each region to determine whether there were morphological differences between core and range.

We used a multi-scale pattern analysis (MSPA) to explore the spatial structure within our data. We implemented a PCA to regress independent variables with predictors of spatial connectivity, using “Adegenet” version 1.4-2 (Jombart et al. 2015) package in R (R Core Team 2013). For our predictors of spatial connectivity we used Moran’s Eigenvector Maps (MEMs), as these have proven to be strong spatial predictors (Jombart et al. 2009). This approach allows for the identification of MEMs in decreasing order of the spatial scale so that MEM1 relates to the broadest spatial scale. We limited the upper bounds of the connection network to enable comparisons between South African and Australian patterns by determining a ceiling value based on both the connection networks that allowed for all locations to be connected at least once to the network. We identified important MEMs based on the inflection point of all the MEMs explanatory power. Using *R*^2^, the determination coefficient of the MSPA, we quantified the association between our independent variables and the MEMs, by strength. We assessed the signals of spatial structure, for morphological and environmental variables, for each region separately, and separately for adult males, adult females, and unsexed juveniles. We conducted a redundancy analysis (RDA) to explain morphology adjusted by environmental variables at all spatial scales. This allowed us to combine both the environmental variables and the spatial predictors to explain the morphological variation detected.

To determine the statistical significance of the correlation between environmental and morphometric traits we created Generalized Additive Mixed Models (GAMMs) for each continent and morphological trait using the environmental factors as explanatory variables. We selected the best models based on the Akaike Information Criterion (AIC) (Akaike 1992). The covariates were tested for significance using backward selection retaining only the significant covariates. In instances where more than one variable was retained we tested for colinearity between them. If two variables showed a high correlation then the model was run separately for each to determine whether significance was maintained for each separate variable. If there was no significance, the variable was removed, if both variables were found to be significant then both were retained. We ran the retained model for each regional dataset and then within each region for adult males, adult females, and unsexed juveniles separately. This allowed us to determine whether the significant variables detected were sex linked. Mann Whitney U tests were carried out for the initial identification of possible spatial sorting patterns using Statistica 12 (Dell Inc. 2015).

## Results

### South Africa

Significant differences were detected between core and range edge starling populations in South Africa for wing load (U= 856, p<0.05, Edge > Core), bill width (U=745, P<0.05, Core > Edge), bill ratio (U=795.5, P<0.05, Edge > Core), head ratio (U=740, P<0.05, Edge > Core) and tarsus ratio (U=594, P<0.05, Edge > Core).

The MSPA of morphological traits showed that the first two axis accounted for 58.89% variation of the entire dataset and, when separated, explained 63.13% for females, 52.28% for males and 54.30% for juveniles. At broader scales bill shape is more discriminating for females primarily but also in males for bill length (MEM2) and juveniles for bill width (MEM1). Wing traits are slightly discriminating for female in wing length (MEM 8) however this effect is more pronounced in males wing load (MEM 23) and wing length (MEM 23-60) (Table S3).

The MSPA of environmental factors showed that the first two axes accounted for 77.61% of the variation in the entire dataset and when separated explained 97.00% for females, 87.29% for males, and 86.77% for juveniles. The environmental data identified MEMs at the broadest spatial scales (Table S4) and as expected these MEMs explained the majority of the variation (between 61% and 97%) in environmental factors. The only exception to this was human impact which explained 20.50% (entire), 17.06% (females), 18.40% (males), and 17.60% (juveniles) of the variation. MEM1 in particular was strongly related to distance from the introduction site, explaining between 73.03% and 97% of the variation across all iterations (Table S4).

When morphological and environmental data were combined via redundancy analysis for all of South Africa, MEMs were identified at the broadest spatial scales and explained a greater degree of the variation in traits (26% to 76%) when compared to morphometric data alone (Table S5). MEM1 was orthogonal to all other MEMs and was linked primarily to bill traits as well as to distance from the introduction site among other environmental traits when the data was assessed in its entirety (Fig. 2A), for females (Fig. 2B) and for juveniles (Fig 2D) whils males showed bill traits explained by MEMs 1 and 2 (Fig. 2C)

**Figure 2:**
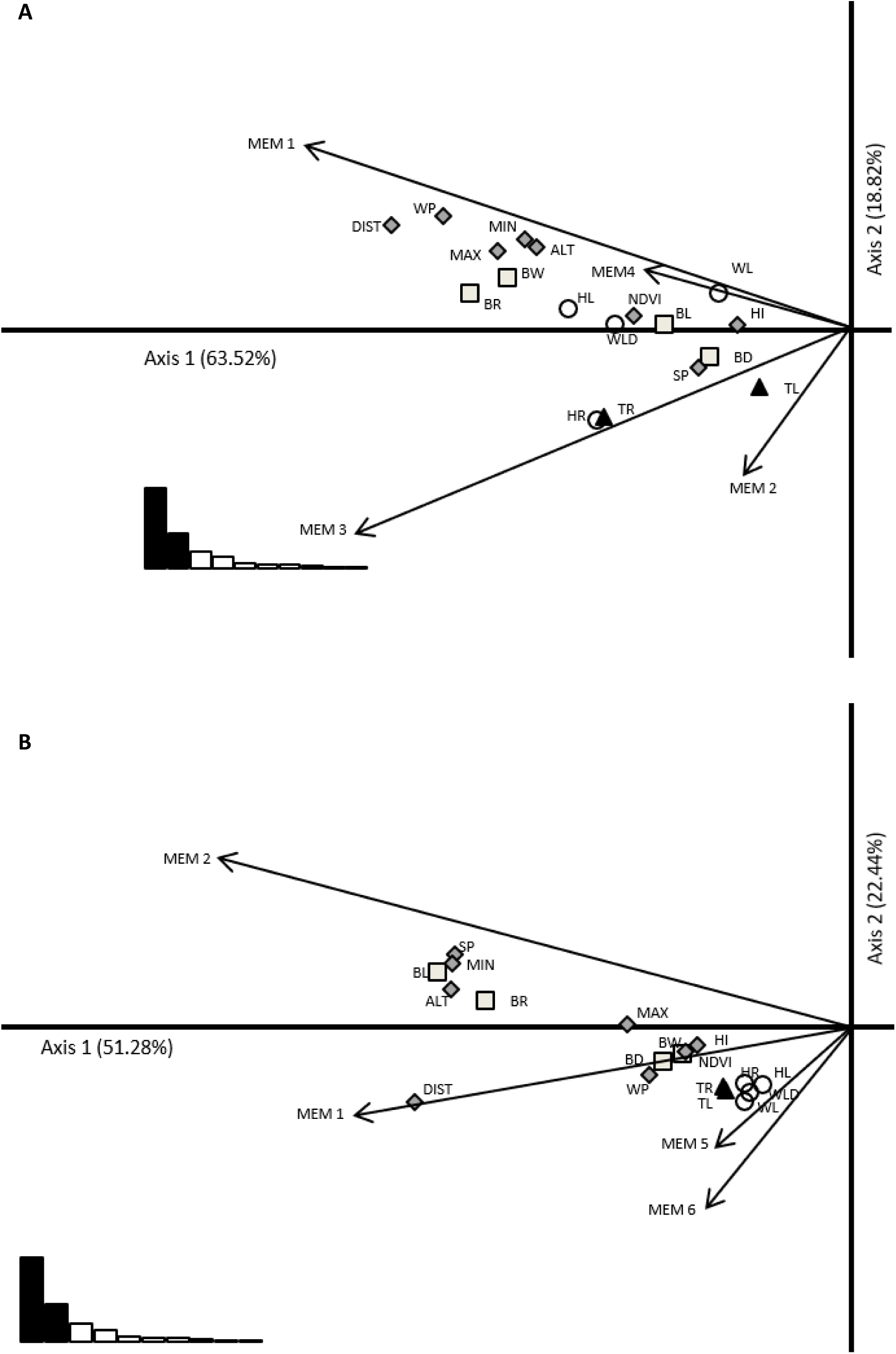
Results of environmental and morphometric analysis using MSPA redundancy analysis for the South African dataset. Eigen values are shown as insets and MEMs indicate predictors of spatial scales. A: Adult Females and B: Adult Males. Filled squares represent traits related to bill morphology, filled triangles traits related to tarsus, open circles traits related to dispersal, and filled diamonds environmental variables. WL: wing length, WLD: wing loading, HL: head length, HR: head ration, BW: bill width, BL: bill length, BD: bill depth, BR: bill ratio, TL: tarsus length, TR: tarsus ratio, DIST: distance from the introduction site, ALT: altitude, MIN: minimum temperature, MAX: maximum temperature, SP: mean summer precipitation, WP: mean winter precipitation, NDVI: normalised difference vegetation index AND HI: human impact.

The GAMM models revealed that sex was significantly linked to tarsus length, tarsus ratio, bill length, bill width, bill ratio, head length and head ratio (Table 2), explaining between 4.33% and 31.76% of the variation across the entire South African dataset.

**Table 2:**
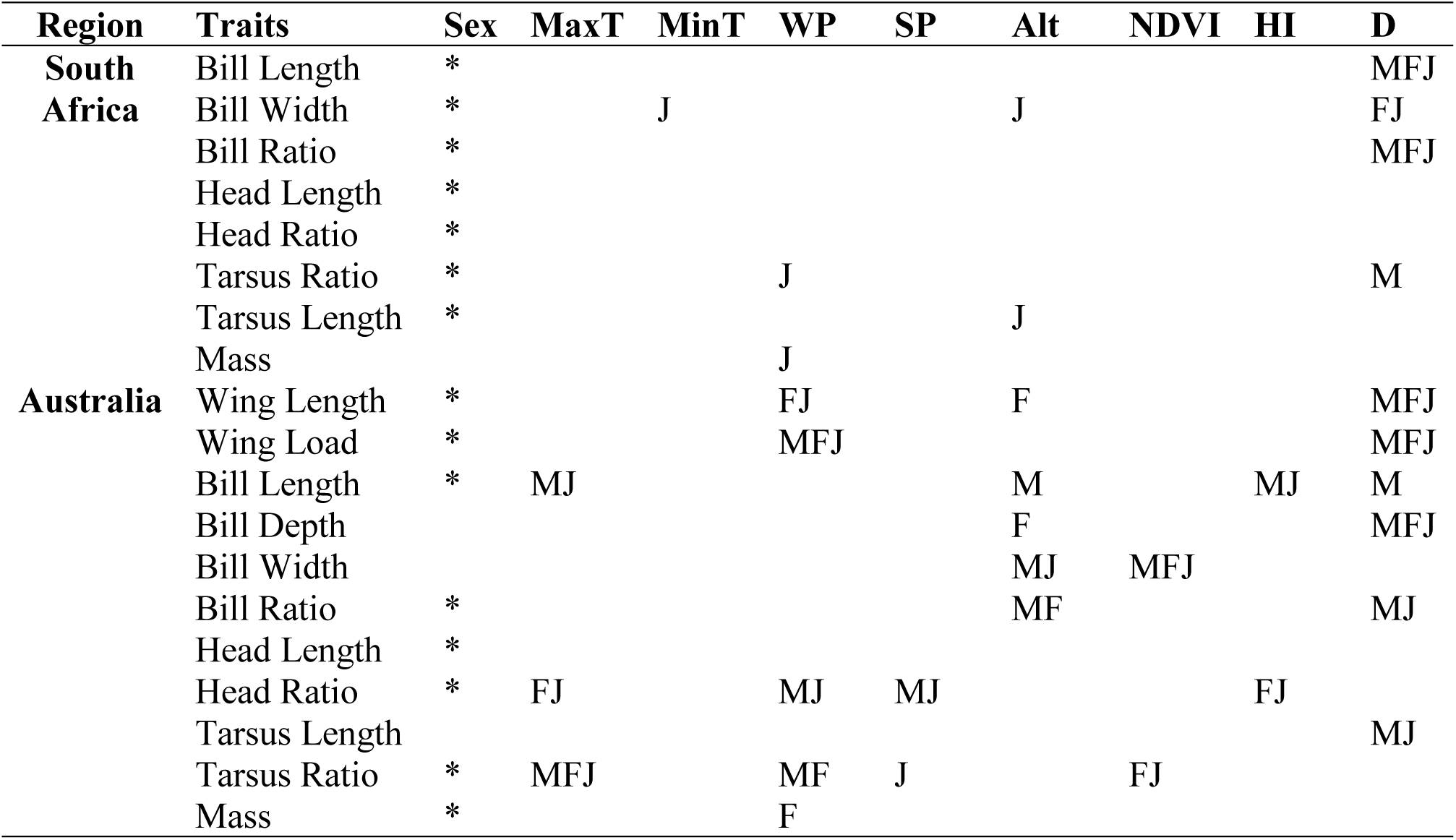
Summary of environmental variables (MaxT: max temperature, MinT: minimum temperarture, WP: Winter precipitation, SP: Summer precipitation,Alt: Altitude, NDVI: Normalised difference vegetation index, HI: Human Impact, D: Distance from introduction site and L: Locality) with significant effect on starling traits measured in this study using generalized additive models and populations in South Africa and Australia. All signs suggest significance, (*) indicate significance of variables when analysed with sex included as a factor, (M: males, F: females and J: juveniles) indicate the significance of factors when sexes were analysed separately.

For all three groups, bill measurements were linked to core distance. Head measurments were sex linked and juveniles showed links to winter precipitation in tarsus ratio and mass whilst linked to altitude in summer precipitation. Males showed a link between tarsus ratio and distance from the introduction site (Table 2).

### Australia

For Australia significant differences between core and range edge were detected in wing load (U= 738, p<0.05, Edge > Core), wing length (U=656, P<0.05, Edge > Core), tarsus length (U= 884, p<0.05, Core > Edge) bill width (U=922, P<0.05, Core > Edge) and bill depth (U=533, P<0.05, Core > Edge).

The MSPA of morphological traits showed that the first two axis accounted for 74.50% of the entire dataset and when separated explained 72.62% for females, 68.61% for males, and 71.38% for juveniles (Table S6). For the Australian dataset MEMs were identified at large spatial scales and no structure was found at fine scales. Wings and BD are the most discirminant for females (MEM 1) and wings and tarsus for males (MEM1)

The MSPA of environmental traits showed that the first two axis accounted for 88.43% of the variation found in the entire dataset and when separated explained 88.16% for females, 87% for males, and 90.20% for juveniles (Table S7). The environmental data identified MEMs at the largest spatial scales, similarly to South Africa these explained the majority of the variation (between 57.99% and 94.24%) in environmental traits. However, human impact, altitude and minimum temperature were explained to a lesser degree by the MEMs ranging between 24.69% and 56.09%. MEM1 in particular was strongly related to distance from the introduction site, explaining between 88.36% and 92.16% of the variation across all iterations

When morphological and environmental data were combined via redundancy analysis for all the Australian data (Fig. 4A, Table S8) MEMs were identified at the broadest scale. MEM1 was orthogonal to the other MEM

s and was related to bill traits but also dispersal traits and strongly related to distance from the introductory site in the entire dataset (Fig. 3A) and in females (Fig 3B). Males dispersal traits and not bill traits were stongly linked to MEM1 (Fig 3C) however juveniles showed a poor relationship between MEM1 and dispersal linke traits (Fig. 3D)

**Figure 3:**
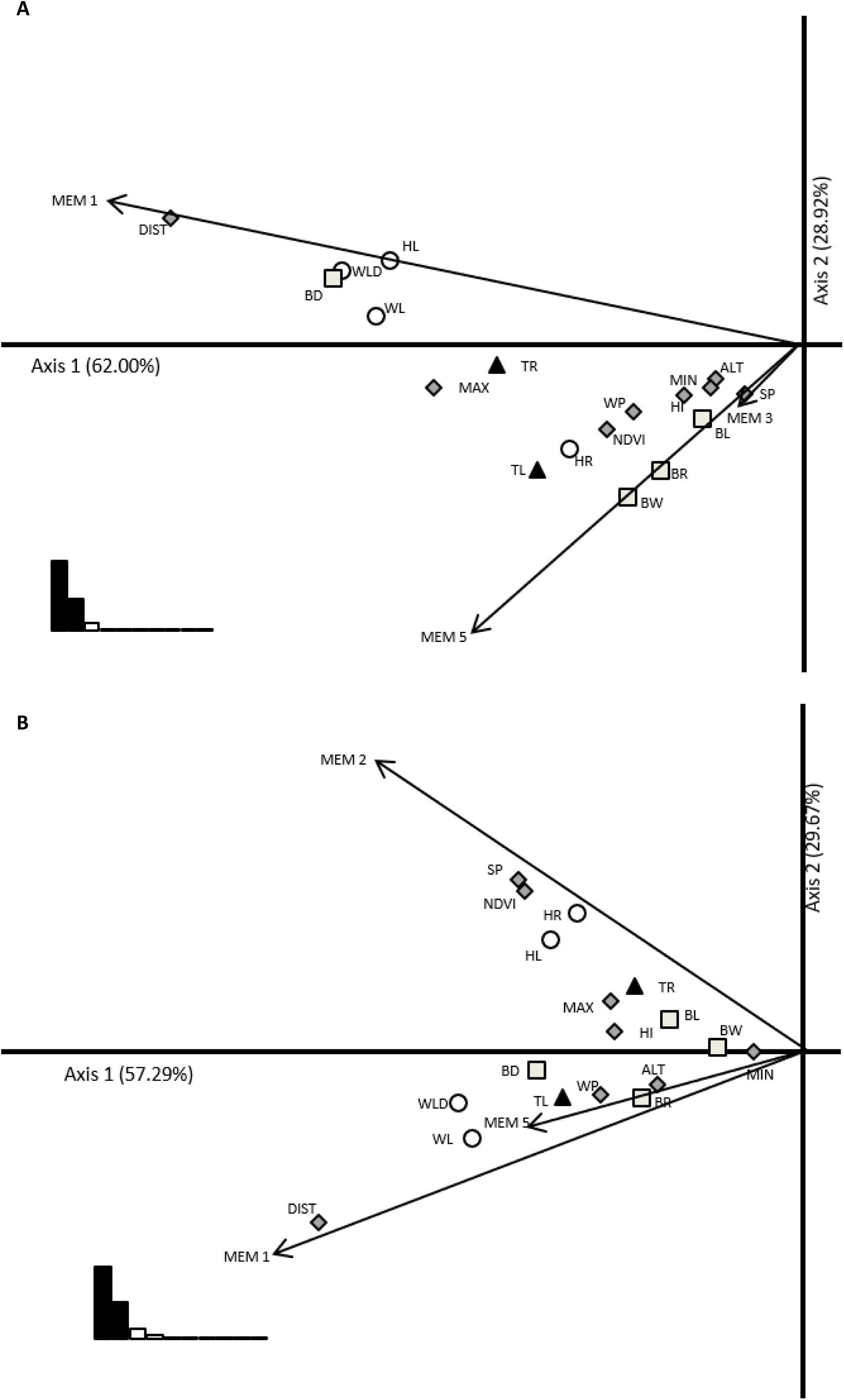
Results of environmental and morphometric analysis using MSPA redundancy analysis for the Australian Dataset. Eigen values are shown as insets, MEMs indicate predictors of spatial scales. A: Adult Females and B: Adult Males. Filled squares represent traits related to bill morphology, filled triangles traits related to tarsus, open circles traits related to dispersal, and filled diamonds environmental variables. WL: wing length, WLD: wing loading, HL: head length, HR: head ration, BW: bill width, BL: bill length, BD: bill depth, BR: bill ratio, TL: tarsus length, TR: tarsus ratio, DIST: distance from the introduction site, ALT: altitude, MIN: minimum temperature, MAX: maximum temperature, SP: mean summer precipitation, WP: mean winter precipitation, NDVI: normalised difference vegetation index AND HI: human impact.

The GAMM models revealed that sex was significantly linked to wing length, wing load, bill length, bill depth, bill ratio, head length, head ratio and tarsus ratio and explained between 6.70% and 29.43% of the variation (Table 2) in the entire Australian dataset. Wing length and load were significantly associated with distance from the introduction site in all three groups, furthermore the three groups showed links between wing load and winter precipitation. In wing length further links were found for juveniles and females to winter precipitation and altitude in females alone.

Tarsus length was significantly linked to distance from the introduction site for males and juveniles. Tarsus ratio was linked to maximum temperature in all groups, males were further linked to winter precipitation, females were linked to winter precipitation and NDVI. Juvenile’s tarsus ratio was linked to summer precipitation and NDVI. Head ratio in juveniles was linked with maximum temperature, winter precipitationm, summer precipitation and human impact. In female’s head ratio was linked to maximum temperature human impact whilst in males it was linked to precipitation, head length merely displayed differentiation between the sexes. Mass was linked to winter precipitation in females. Bill length was associated with maximum temperature, distance, human impact and altitude in males. Juveniles bill length was linked to human impact and maximum temperature. Bill depth was linked to distance from introduction site in all groups and further to altitude in females. Bill width was linked to NDVI in all groups and to altidude in males and juveniles. Bill ratio was linked to altitude and distance from introduction site for males whilst females were linked to altitude and juveniles to distance from introduction site (Table 2).

## Discussion

### Dispersal Traits

Theory predicts that species experiencing spatial sorting should display enhanced dispersability and its associated traits as distance from the original distribution or, for invasive species, initial introduction site, increases (Shine et al. 2011). In Australia, starlings presented strong proof of spatial sorting with both wing length and wing loading being strongly linked to distance from the original site of introduction (see Table 2, and Fig. 3). In contrast, starlings in South Africa did not show signs of spatial sorting of dispersal traits, but instead for traits associated with intraspecific competition such as bill length, width and ratio (see Table 2, and Fig. 2). The patterns detected in Australia match findings in other invasive species like the cane toads in Australia (Phillips et al. 2006), Indian mynas in South Africa (Berthouly-Salazar et al. 2012), pygmy grasshoppers in Sweden (Berggren et al. 2012), and ladybirds in Europe (Lombaert et al. 2014). The South African findings were not unexpected as it is not the first study to find no patterns of spatial sorting of wing traits in *S. vulgaris* (Bitton and Graham 2015). Similar to our findings for South African starlings, North American starling invasions show no evidence for increased wing length along the invasion path which was attributed to potential long distance dispersal by adults (Bitton and Graham 2015).

The apparent differences between South Africa and Australia found here could be attributed to differences in population connectivity, sampling strategies, or local conditions; however, there is strong support for the hypothesis of population connectivity. Previous genetic analyses of starlings in Western Australia found clear population differentiation between core and range edge populations (Rollins et al. 2009, Rollins et al. 2011) and no evidence for genetic bottlenecks, suggesting the existence of distinct genetic populations with limited geneflow. A lack of gene flow may therefore prohibit the dilution of any accrued morphological differences due to spatial sorting. In contrast, similar genetic analyses of South African starlings (Berthouly-Salazar et al. 2013) found strong genetic connectivity between core and range edge populations as well as low IBD (isolation by distance), suggesting frequent extreme long distance dispersal (LDD) events. The latter is in agreement with dispersal patterns from across the native range of starlings (Hui et al. 2012). Despite spatial sorting increasing the number of strong dispersers at the range edge, frequent extreme LDD events are expected to dilute patterns of spatial sorting by introducing less effective dispersers from the core population into the range edge. Klein et al. (2006) showed that a fat-tailed dispersal kernel, indicative of frequent LDD, results in an overall panmictic population. The likelihood of extreme LDD events occurring may further be related to density- and habitat-dependant dispersal. Extreme LDD events are expected to occur more readily in poorer habitats, as a spreading population is expected to follow the “good stay, bad disperse” strategy for spread (Hui et al. 2012). One such example is the densely populated range core, where intense competition for available resources could trigger extreme LDD events (Muller-Landau et al. 2003).

While starlings are migratory in their native range (Thomson 1921) they are generally non-migratory in invaded regions (Dolbeer 1982). Starlings have been noted to migrate in some non-native regions (Kessel 1953; Berthouly-Salazar et al. 2013), suggesting that dispersal rates may change when disperser encounter changing environmental conditions, whether this is a result of favourable or unfavourable conditions requires further assessment. Berthouly-Salazar et al. (2013) indicated that accelerated dispersal of starlings in South Africa is linked to increased contact with changing precipitation regimes, a finding that is supported by ecological models (Hui et al. 2012). It is likely that individuals from the range core dispersing outward encounter regions of changing environmental conditions and, by following the “good stay, bad disperse” rule (Hui et al. 2012), increase their dispersal rate, resulting in poor dispersers obscuring the effects of spatial sorting towards the range edge.

Alternate factors that may explain the differences of patterns detected between South African and Australian starling populations may include differences in sampling, clinal variation, environmental heterogeneity or even differences in dispersal rates and demography. The maximum distance from introduction sites were comparable between South Africa and Australia as was the sampling effort employed. Clinal variation may lead to changes in traits across latitudinal ranges. Specifically, variation in body size may change depending on geographic location (Graves 1991) or environmental traits (Hamilton 1958, 1961; James 1970). Starlings in Australia have been shown to conform to Bergmann’s (Bergmann 1847) and Allen’s (Allen 1877) rule suggesting that starlings may undergo rapid adaptive change in relation to clinal variation (Cardilini et al. 2016).

Demographic differences such as introduction history and propagule pressure may also play a role in the patterns detected; to date we know of only one introduction event into South Africa of 18 individuals in 1897, Australia however has records of multiple introductions and releases accounting for more than 200 individuals (Long 1981). This difference in propagule pressure resulted in differences in genetic variation and may have affected dispersal and lead to starlings in Australia not experiencing an initial lag phase of dispersal or harbouring more genetically based phenotypic variation in traits related to dispersal. Dispersal rates may also have differed between the two introductions, Hui et al. (2012) found that in South Africa starlings initially dispersed slowly during the establishment phase (6.1 km/yr) before increasing their dispersal rate to 25.7 km/yr. We estimated that Australian starlings may have experienced an average dispersal rate of 20.7 km/yr, ranging between 17.3 to 40.4 km/yr. Finally this study did not account for biotic differences, such as predation or competition with local species, which may play a role in the differences in spatial sorting of traits between South African and Australia.

Lastly, we found Australian starlings to show a significant positive association between distance from introduction site and wing length across all sexes and age classes. This was unexpected, as the majority of dispersal events in starlings are usually by juveniles (Kessel 1957). We also found juvenile wing length to differ significantly from adult males and females (see Table S2) and thus expected a more pronounced effect in juveniles. This suggests that dispersal trait changes are lifelong and not age-specific.

## Foraging Traits

Literature suggests that the increased selection on dispersal-linked traits may likely lead to evolutionary trade-offs such as decreased reproductive potential (Hughes et al. 2003). For example, the grasshopper, *Tetrix subulata,* has two distinct morphs; a highly dispersive long winged morph and a short winged morph with increased reproductive ability (Steenman et al. 2014). Similarly, it has been recently shown that invasive cane toads in the edge of their invasive range show increased dipersibility but decreased reproductive output (Hudson et al. 2015). Other trade-offs include competitive ability, with theoretical models predicting that these are strongly selected against at the range front (Burton et al. 2010). Limited empirical evidence exists for this, and our study is one of the first to find competitive and foraging traits related to distance from introduction site.

Bill length and bill ratio in South Africa, were significantly linked to distance from the introduction site, as was bill depth in Australia. It has further been suggested that if the costs related to the production and maintenance of augmented dispersal are off-set sufficiently by increased mobility and foraging capability, trade-offs such as competitive versus dispersal traits may disappear (Berggren et al. 2012). If selection is driving these trade-offs it may result in reduced competition with conspecifics at the invasion front, increasing resource availability and fitness, as was detected in cane toads (Brown et al. 2013). Alternatively, natural selection could act at the range front and select for phenotypes that increase reproductive output at low population densities (Holt et al. 2005; Perkins et al. 2013). The above mentioned selective forces may have played a role in directing the observed relationship between bill traits and distance from introduction site for invasive starlings in Australia and South Africa. However, we cannot discount that these observations may have resulted from environmental variation (phenotypic plasticity) or local competition (Radford and Du Plessis 2003; Grant and Grant 2006) as climate variation has been shown to significantly affect starling bill surface area (Cardilini et al. 2016).

This study did not account for potential biotic interactions in shaping the spatial distribution of traits and may therefore have not accounted for the possibility that invasive starlings may have experienced different levels of predation and competition in South Africa and Australia. A cursory assessment of raptor diversity, based on the global raptor information network (Rosenberry 2015), shows that South Africa has almost three times the raptor diversity of Australia. Furthermore, the pressure exerted from predation is strongly linked to the density of generic raptors and as such is expected to be greater in South Africa. This difference in numbers of possible predators may explain why South African starling show increased competitive traits with an increased distance from the core. It may be possible that interspecific interaction may play a more important role than spatial sorting in moulding morphology, although further research is required to support this notion.

## Environmental Filters

Environmental heterogeneity may also play a role in preventing or facilitating spatial sorting patterns. South Africa, and especially the Western Cape province, is an extremely heterogeneous landscape (Goldblatt and Manning 2002). This heterogeneous environment may have lead to more frequent dispersal events in stralings as habitat suitability may be highly variable, confounding spatial sorting patterns. This is further strengthened by the lack of interactions between South African starlings morphological traits and environmental factors. Australian starlings however showed more linkages between environmental factors and morphological traits, which may be a result of the sampling design occurring across a strong environmental cline.

Furthermore, a closer look at other invasive bird species shared between South Africa and Australia is warranted to determine if the patterns detected in this study are a general occurrence or specific to starlings. Differences in the introduction point and avenues of dispersal may potentially play a role in affectiong patterns of spatial sorting. South African starlings were introduced into the southwest of the region and optimal conditions occurred more readily to the east (Hui et al. 2012), encouraging longitudinal spread, whilst Australian starling were introduced in the south east and primarily spread latidudinally. Within the same study system it is worth including a wider range of traits, as this study was limited to the overlap of two separate datasets. A concerted sampling and measuring effort across both continents could allow for more traits related to dispersal to be included such as tail length or graduation (Dawideit et al. 2009), wing pointedness (Baldwin et al. 2010) and wing to tail ratio which all provide stronger measures of manoeuvrability (e.g. Bitton and Graham 2015). Measures of these traits, reproductive traits and more competitive traits could allow for empirical tests of the many theorised patterns resulting from spatial sorting, such as trade-offs between dispersal and reproduction (Hughes et al. 2003) or competition (Burton et al. 2010).

## Conclusion

This study provides evidence that starlings are experiencing spatial sorting in Australia and South Africa but for different traits in each country. Within Australia dispersal linked traits, wing load and wing length, were shown to correlate with distance from the introduction site suggesting classic spatial sorting. In contrast, patterns of spatial sorting were not detected in the dispersal traits of South African starlings; instead traits related to the resource acquisition, bill length and bill ratio, were associated with distance from the introduction site. These findings are likely a result of the alternate selective pressures between the countries. Australian starlings appear to be steadily expanding their range through sequential founding as suggested by genetic studies (Rollins et al. 2009, 2011), which would favour spatial sorting of dispersal traits. However in South Africa starlings are experiencing frequent long distance dispersal events as suggested by genetic studies (Berthouly-Salazar et al. 2013), these events may potentially erode spatial sorting patterns on dispersal-linked traits. Despite our findings of a lack of spatial sorting on dispersal linked traits, SA starlings still show patterns of increased rates of spread (Hui et al. 2012), suggesting that frequent extreme LDD events may dilute patterns of spatial sorting through the translocation of starlings from the core to the range edge. Furthermore, this study identified a relationship between resource acquisition traits and distance from the introduction site. This pattern was more pronounced in South Africa, which is likely a result of the highly heterogeneous environment characterising the area invaded by starlings. These findings demonstrate the importance of assessing ecological patterns within common species across continental scales, as it may help to identify potential factors affecting range expansions.

## Acknowledgements

We would like to thank Nikki Phair for providing insights into the manuscript. DJP received a MSc Scholarship from the National Research Foundation (NRF) of South Africa. CH is supported by the South African Research Chair Initiative (SARChI), the National Research Foundation of South Africa (grant nos. 81825 and 109244), and the Australian Research Council (Discovery Project DP150103017). JLR acknowledges funding from the National Research Foundation of South Africa (grant no. 91117) and Stellenbosch University’s Subcommittee B. VV was funded by the National Research Foundation, South Africa (grant 85265) and SANBI’s Invasive Species Programme. The South African collections protocols were approved by, The animal care committee of Stellenbosch University (N°10NP_CEC01) and CapeNature (N°AAA-004-00499-0035).

## Supplementary Information

**Table S1:** Morphological traits measured in invasive populations of S*. vulgaris* in Australia and South Africa

| Trait | Measurement Details: |
| --- | --- |
| Tarsus length | Distance from the tibio-tarsus joint to the distal end of the tarso-metatarsus |
| Wing length | Distance from the carpal joint to the tip of the longest primary |
| Bill length | Distance from the bill tip to the skull at nasal-frontal hinge |
| Bill width | Distance at the proximate edge of the nostrils |
| Bill depth | Distance at the proximate edge of the nostrils |
| Head length | Distance from the tip of bill to the back of head, minus bill length |
| Weight |  |

**Table S2:**
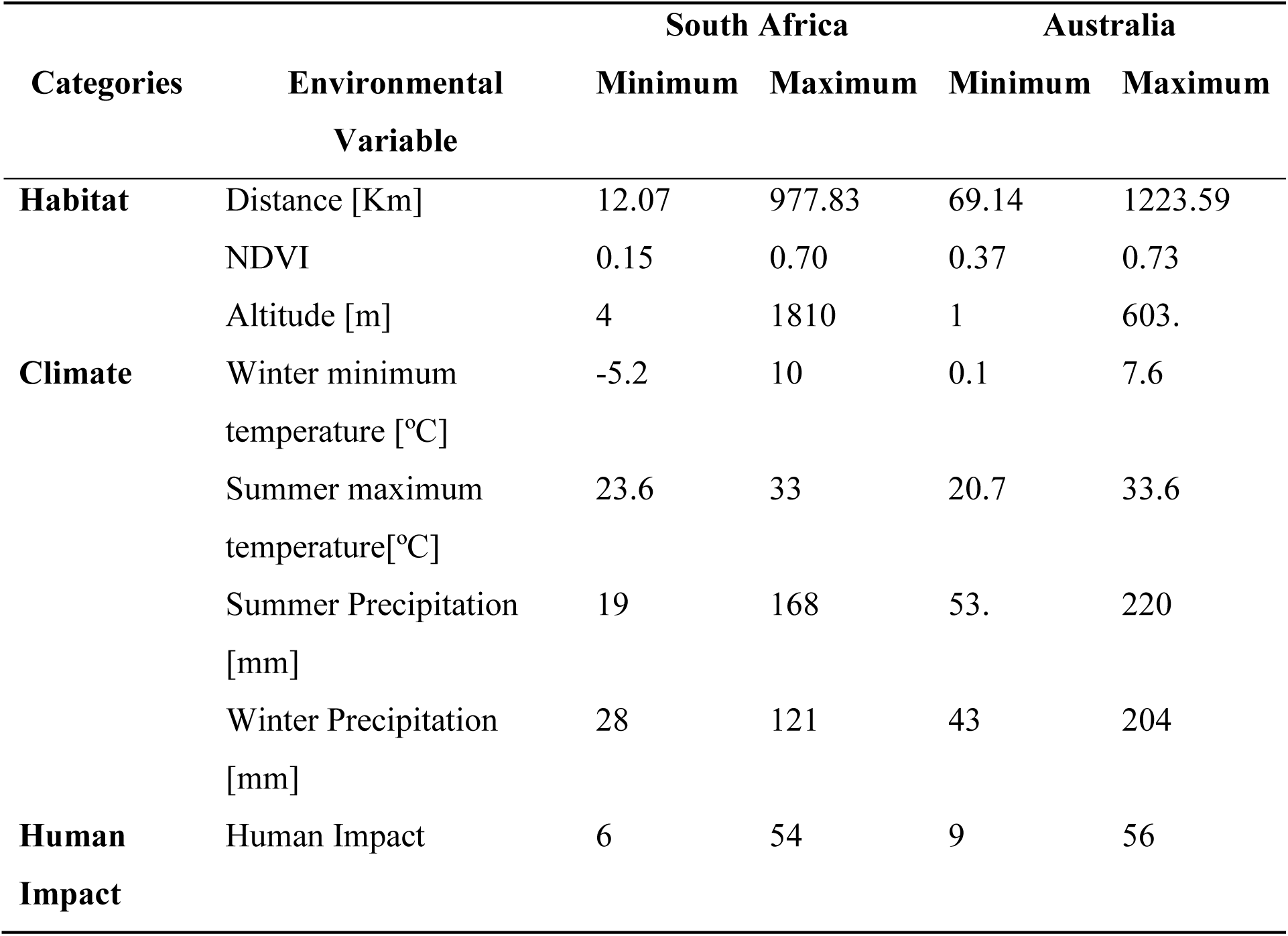
Summary of environmental conditions within study regions

**Table S3:** Summary of important MEM R2 values for starling morphological traits within the South African dataset.

| Dataset | MEM | Wing Load | Wing Length | Tarsus Length | Bill Length | Bill Depth | Bill Width | Head Length | Bill Ratio | Head Ratio | TR |
| --- | --- | --- | --- | --- | --- | --- | --- | --- | --- | --- | --- |
| All | 1 | 0,011 | 0,005 | 0,000 | 0,026 | 0,007 | 0,094 | 0,003 | 0,102 | 0,053 | 0,150 |
|  | 28 | 0,042 | 0,040 | 0,001 | 0,002 | 0,000 | 0,000 | 0,004 | 0,002 | 0,004 | 0,001 |
|  | 71 | 0,045 | 0,043 | 0,000 | 0,002 | 0,001 | 0,002 | 0,004 | 0,001 | 0,009 | 0,004 |
|  | 92 | 0,065 | 0,054 | 0,000 | 0,004 | 0,011 | 0,002 | 0,003 | 0,000 | 0,001 | 0,000 |
|  | 29 | 0,046 | 0,040 | 0,005 | 0,006 | 0,002 | 0,004 | 0,009 | 0,009 | 0,010 | 0,002 |
|  | Total | 0,209 | 0,182 | 0,006 | 0,039 | 0,021 | 0,101 | 0,023 | 0,114 | 0,077 | 0,157 |
| Females | 1 | 0,068 | 0,059 | 0,000 | 0,090 | 0,018 | 0,205 | 0,091 | 0,225 | 0,006 | 0,027 |
|  | 3 | 0,034 | 0,001 | 0,001 | 0,118 | 0,012 | 0,100 | 0,113 | 0,171 | 0,051 | 0,102 |
|  | 4 | 0,015 | 0,038 | 0,035 | 0,036 | 0,001 | 0,144 | 0,004 | 0,131 | 0,022 | 0,001 |
|  | 7 | 0,061 | 0,078 | 0,064 | 0,000 | 0,002 | 0,034 | 0,141 | 0,011 | 0,003 | 0,003 |
|  | 8 | 0,077 | 0,115 | 0,022 | 0,006 | 0,008 | 0,011 | 0,004 | 0,000 | 0,018 | 0,001 |
|  | 23 | 0,023 | 0,043 | 0,007 | 0,001 | 0,179 | 0,000 | 0,007 | 0,000 | 0,018 | 0,007 |
|  | 28 | 0,041 | 0,004 | 0,051 | 0,060 | 0,087 | 0,001 | 0,016 | 0,013 | 0,076 | 0,095 |
|  | 29 | 0,141 | 0,043 | 0,017 | 0,021 | 0,004 | 0,035 | 0,005 | 0,007 | 0,047 | 0,073 |
|  | 31 | 0,036 | 0,143 | 0,006 | 0,019 | 0,029 | 0,014 | 0,047 | 0,012 | 0,050 | 0,063 |
|  | Total | 0,496 | 0,525 | 0,202 | 0,352 | 0,340 | 0,544 | 0,428 | 0,571 | 0,292 | 0,372 |
| Males | 2 | 0,001 | 0,000 | 0,002 | 0,177 | 0,010 | 0,004 | 0,002 | 0,070 | 0,002 | 0,000 |
|  | 20 | 0,065 | 0,025 | 0,095 | 0,047 | 0,039 | 0,046 | 0,000 | 0,080 | 0,002 | 0,016 |
|  | 21 | 0,073 | 0,039 | 0,025 | 0,009 | 0,003 | 0,001 | 0,010 | 0,001 | 0,072 | 0,059 |
|  | 23 | 0,141 | 0,242 | 0,002 | 0,002 | 0,036 | 0,019 | 0,002 | 0,018 | 0,009 | 0,072 |
|  | 60 | 0,088 | 0,112 | 0,024 | 0,002 | 0,014 | 0,018 | 0,014 | 0,011 | 0,005 | 0,014 |
|  | Total | 0,368 | 0,418 | 0,148 | 0,237 | 0,102 | 0,087 | 0,029 | 0,180 | 0,089 | 0,161 |
| Juveniles | 1 | 0,013 | 0,014 | 0,013 | 0,009 | 0,027 | 0,116 | 0,003 | 0,098 | 0,040 | 0,087 |
|  | 2 | 0,006 | 0,004 | 0,087 | 0,095 | 0,006 | 0,015 | 0,058 | 0,086 | 0,060 | 0,048 |
|  | 5 | 0,001 | 0,002 | 0,026 | 0,038 | 0,018 | 0,004 | 0,057 | 0,030 | 0,068 | 0,002 |
|  | 45 | 0,067 | 0,066 | 0,004 | 0,001 | 0,000 | 0,044 | 0,002 | 0,011 | 0,011 | 0,018 |
|  | 78 | 0,015 | 0,009 | 0,001 | 0,001 | 0,000 | 0,084 | 0,004 | 0,035 | 0,001 | 0,003 |
|  | 87 | 0,002 | 0,004 | 0,002 | 0,032 | 0,005 | 0,002 | 0,151 | 0,018 | 0,045 | 0,002 |
|  | Total | 0,104 | 0,099 | 0,132 | 0,176 | 0,057 | 0,266 | 0,274 | 0,277 | 0,225 | 0,161 |

Highlighted cells represent cases where greater than 10% of the variation in a trait is explained by a MEM

**Table S4:** Summary of important MEM *R^2^* values for environmental conditions in South Africa.

| Dataset | MEM | HI | Winter<br>prec | Summer<br>prec | alt | Max temp | Min temp | NDVI | DISTANCE |
| --- | --- | --- | --- | --- | --- | --- | --- | --- | --- |
| All | 1 | 0,014 | 0,103 | 0,032 | 0,383 | 0,004 | 0,174 | 0,028 | 0,949 |
|  | 2 | 0,038 | 0,425 | 0,170 | 0,019 | 0,014 | 0,002 | 0,035 | 0,006 |
|  | 3 | 0,011 | 0,023 | 0,127 | 0,097 | 0,255 | 0,128 | 0,093 | 0,004 |
|  | 4 | 0,006 | 0,124 | 0,129 | 0,003 | 0,076 | 0,004 | 0,170 | 0,007 |
|  | 5 | 0,005 | 0,015 | 0,102 | 0,280 | 0,076 | 0,395 | 0,157 | 0,003 |
|  | 6 | 0,044 | 0,055 | 0,010 | 0,001 | 0,255 | 0,008 | 0,093 | 0,000 |
|  | 9 | 0,089 | 0,003 | 0,137 | 0,066 | 0,004 | 0,041 | 0,049 | 0,000 |
|  | Total | 0,205 | 0,748 | 0,707 | 0,851 | 0,684 | 0,753 | 0,625 | 0,970 |
| Females | 1 | 0,106 | 0,676 | 0,001 | 0,459 | 0,583 | 0,535 | 0,155 | 0,676 |
|  | 2 | 0,033 | 0,044 | 0,417 | 0,001 | 0,215 | 0,032 | 0,209 | 0,002 |
|  | 4 | 0,031 | 0,006 | 0,421 | 0,116 | 0,007 | 0,012 | 0,356 | 0,053 |
|  | Total | 0,171 | 0,726 | 0,840 | 0,577 | 0,805 | 0,579 | 0,720 | 0,730 |
| Males | 1 | 0,036 | 0,120 | 0,048 | 0,202 | 0,004 | 0,079 | 0,029 | 0,856 |
|  | 2 | 0,146 | 0,131 | 0,559 | 0,426 | 0,282 | 0,521 | 0,158 | 0,009 |
|  | 3 | 0,000 | 0,029 | 0,002 | 0,205 | 0,018 | 0,200 | 0,201 | 0,040 |
|  | 4 | 0,002 | 0,024 | 0,111 | 0,005 | 0,200 | 0,043 | 0,271 | 0,020 |
|  | Total | 0,184 | 0,304 | 0,720 | 0,837 | 0,504 | 0,844 | 0,659 | 0,925 |
| Juveniles | 1 | 0,010 | 0,229 | 0,000 | 0,569 | 0,062 | 0,359 | 0,118 | 0,900 |
|  | 2 | 0,070 | 0,373 | 0,079 | 0,050 | 0,144 | 0,010 | 0,017 | 0,028 |
|  | 3 | 0,036 | 0,082 | 0,573 | 0,001 | 0,212 | 0,000 | 0,341 | 0,000 |
|  | 5 | 0,006 | 0,011 | 0,048 | 0,014 | 0,277 | 0,037 | 0,088 | 0,002 |
|  | 7 | 0,053 | 0,005 | 0,053 | 0,181 | 0,058 | 0,212 | 0,238 | 0,000 |
|  | Total | 0,176 | 0,699 | 0,754 | 0,815 | 0,753 | 0,617 | 0,802 | 0,930 |
| Highlighted cells represent cases where greater than 10% of the variation in a trait is explained by a MEM |  |  |  |  |  |  |  |  |  |

**Table S5:** Summary of important MEM R2 values of starling morphological traits adjusted by environmental traits in South Africa.

| Dataset | MEM | wingLoad | RwingL | RlegL | RbillL | RbillD | RbillW | RheadL | BR | HR | TR |
| --- | --- | --- | --- | --- | --- | --- | --- | --- | --- | --- | --- |
| All | 1 | 0,245 | 0,101 | 0,008 | 0,246 | 0,171 | 0,564 | 0,098 | 0,536 | 0,441 | 0,690 |
|  | 3 | 0,093 | 0,074 | 0,259 | 0,057 | 0,024 | 0,005 | 0,021 | 0,017 | 0,019 | 0,001 |
|  | 5 | 0,000 | 0,014 | 0,130 | 0,042 | 0,073 | 0,002 | 0,073 | 0,004 | 0,102 | 0,014 |
|  | 6 | 0,024 | 0,013 | 0,214 | 0,074 | 0,040 | 0,049 | 0,173 | 0,084 | 0,003 | 0,060 |
|  | 7 | 0,039 | 0,060 | 0,009 | 0,082 | 0,055 | 0,021 | 0,146 | 0,042 | 0,046 | 0,001 |
|  | Total | 0,402 | 0,262 | 0,620 | 0,502 | 0,363 | 0,641 | 0,511 | 0,683 | 0,610 | 0,766 |
| Females | 1 | 0,220 | 0,149 | 0,000 | 0,157 | 0,057 | 0,391 | 0,324 | 0,396 | 0,039 | 0,080 |
|  | 2 | 0,051 | 0,005 | 0,360 | 0,010 | 0,014 | 0,021 | 0,071 | 0,025 | 0,064 | 0,177 |
|  | 3 | 0,168 | 0,012 | 0,052 | 0,131 | 0,174 | 0,163 | 0,161 | 0,235 | 0,419 | 0,351 |
|  | 4 | 0,071 | 0,117 | 0,078 | 0,085 | 0,062 | 0,194 | 0,044 | 0,191 | 0,043 | 0,000 |
|  | 7 | 0,039 | 0,060 | 0,009 | 0,082 | 0,055 | 0,021 | 0,146 | 0,042 | 0,046 | 0,001 |
|  | Total | 0,549 | 0,343 | 0,499 | 0,465 | 0,362 | 0,789 | 0,747 | 0,890 | 0,611 | 0,609 |
| Males | 1 | 0,067 | 0,089 | 0,147 | 0,149 | 0,271 | 0,164 | 0,031 | 0,271 | 0,078 | 0,141 |
|  | 2 | 0,006 | 0,000 | 0,003 | 0,498 | 0,051 | 0,086 | 0,022 | 0,343 | 0,029 | 0,014 |
|  | 5 | 0,098 | 0,008 | 0,163 | 0,060 | 0,000 | 0,037 | 0,205 | 0,001 | 0,072 | 0,093 |
|  | 6 | 0,138 | 0,280 | 0,137 | 0,005 | 0,035 | 0,080 | 0,003 | 0,016 | 0,077 | 0,115 |
|  | Total | 0,309 | 0,377 | 0,450 | 0,712 | 0,358 | 0,367 | 0,261 | 0,630 | 0,256 | 0,363 |
| Juveniles | 1 | 0,088 | 0,107 | 0,056 | 0,085 | 0,170 | 0,508 | 0,041 | 0,433 | 0,273 | 0,399 |
|  | 2 | 0,022 | 0,016 | 0,406 | 0,333 | 0,087 | 0,034 | 0,338 | 0,163 | 0,134 | 0,115 |
|  | 7 | 0,003 | 0,018 | 0,075 | 0,019 | 0,387 | 0,001 | 0,062 | 0,009 | 0,019 | 0,041 |
|  | 3 | 0,188 | 0,211 | 0,040 | 0,005 | 0,013 | 0,001 | 0,036 | 0,001 | 0,112 | 0,099 |
|  | Total | 0,301 | 0,351 | 0,576 | 0,442 | 0,656 | 0,544 | 0,477 | 0,607 | 0,538 | 0,654 |

Highlighted cells represent cases where greater than 10% of the variation in a trait is explained by a MEM

**Table S6:** Summary of important MEM *R^2^* values for starling morphological traits in Australia.

| Dataset | MEM | wingLoad | RwingL | RlegL | RbillL | RbillD | RbillW | RheadL | BR | HR | TR |
| --- | --- | --- | --- | --- | --- | --- | --- | --- | --- | --- | --- |
| All | 1 | 0,090 | 0,095 | 0,039 | 0,087 | 0,153 | 0,002 | 0,000 | 0,023 | 0,004 | 0,000 |
|  | 5 | 0,020 | 0,041 | 0,005 | 0,000 | 0,000 | 0,003 | 0,000 | 0,001 | 0,073 | 0,085 |
|  | 6 | 0,027 | 0,041 | 0,001 | 0,032 | 0,002 | 0,000 | 0,001 | 0,014 | 0,033 | 0,072 |
|  | Total | 0,137 | 0,178 | 0,046 | 0,119 | 0,155 | 0,005 | 0,002 | 0,038 | 0,110 | 0,158 |
| Females | 1 | 0,208 | 0,207 | 0,006 | 0,010 | 0,171 | 0,000 | 0,035 | 0,004 | 0,025 | 0,066 |
|  | 2 | 0,006 | 0,001 | 0,007 | 0,021 | 0,015 | 0,037 | 0,000 | 0,042 | 0,031 | 0,051 |
|  | 3 | 0,003 | 0,020 | 0,001 | 0,127 | 0,014 | 0,001 | 0,003 | 0,046 | 0,000 | 0,000 |
|  | 5 | 0,010 | 0,049 | 0,048 | 0,026 | 0,016 | 0,055 | 0,001 | 0,062 | 0,106 | 0,080 |
|  | 13 | 0,006 | 0,003 | 0,021 | 0,026 | 0,030 | 0,000 | 0,099 | 0,010 | 0,046 | 0,012 |
|  | Total | 0,233 | 0,280 | 0,082 | 0,210 | 0,247 | 0,093 | 0,137 | 0,163 | 0,208 | 0,210 |
| Males | 1 | 0,152 | 0,180 | 0,124 | 0,013 | 0,091 | 0,037 | 0,014 | 0,076 | 0,005 | 0,020 |
|  | 2 | 0,036 | 0,012 | 0,013 | 0,106 | 0,052 | 0,035 | 0,089 | 0,002 | 0,183 | 0,112 |
|  | 5 | 0,095 | 0,178 | 0,021 | 0,004 | 0,144 | 0,001 | 0,003 | 0,006 | 0,001 | 0,033 |
|  | 12 | 0,069 | 0,033 | 0,015 | 0,018 | 0,003 | 0,106 | 0,005 | 0,055 | 0,051 | 0,113 |
|  | 14 | 0,013 | 0,017 | 0,002 | 0,172 | 0,016 | 0,002 | 0,078 | 0,071 | 0,111 | 0,044 |
|  | Total | 0,365 | 0,421 | 0,175 | 0,314 | 0,306 | 0,179 | 0,189 | 0,211 | 0,351 | 0,323 |
| Juveniles | 1 | 0,001 | 0,013 | 0,043 | 0,223 | 0,175 | 0,011 | 0,009 | 0,117 | 0,003 | 0,000 |
|  | 5 | 0,028 | 0,009 | 0,002 | 0,014 | 0,002 | 0,000 | 0,003 | 0,006 | 0,039 | 0,068 |
|  | 7 | 0,053 | 0,073 | 0,003 | 0,006 | 0,000 | 0,007 | 0,010 | 0,011 | 0,056 | 0,077 |
|  | Total | 0,082 | 0,096 | 0,049 | 0,242 | 0,177 | 0,019 | 0,022 | 0,133 | 0,097 | 0,145 |
| Highlighted cells represent cases where greater than 10% of the variation in a trait is explained by a MEM |  |  |  |  |  |  |  |  |  |  |  |

**Table S7:** Summary of important MEM *R^2^* values for environmental conditions in Australia.

| Dataset | MEM | HI | winterprec | summerprec | alt | maxtemp | mintemp | NDVI | DISTANCE |
| --- | --- | --- | --- | --- | --- | --- | --- | --- | --- |
| All | 1 | 0,110 | 0,140 | 0,014 | 0,048 | 0,374 | 0,036 | 0,076 | 0,884 |
|  | 2 | 0,138 | 0,200 | 0,587 | 0,214 | 0,183 | 0,158 | 0,552 | 0,010 |
|  | 3 | 0,002 | 0,309 | 0,008 | 0,082 | 0,116 | 0,279 | 0,024 | 0,022 |
|  | 5 | 0,243 | 0,013 | 0,039 | 0,003 | 0,120 | 0,007 | 0,127 | 0,014 |
|  | 6 | 0,008 | 0,078 | 0,110 | 0,083 | 0,098 | 0,056 | 0,037 | 0,002 |
|  | Total | 0,501 | 0,741 | 0,758 | 0,431 | 0,890 | 0,537 | 0,816 | 0,932 |
| Females | 1 | 0,068 | 0,109 | 0,001 | 0,058 | 0,368 | 0,047 | 0,122 | 0,914 |
|  | 3 | 0,140 | 0,167 | 0,513 | 0,177 | 0,056 | 0,104 | 0,381 | 0,007 |
|  | 4 | 0,121 | 0,107 | 0,015 | 0,191 | 0,017 | 0,195 | 0,000 | 0,005 |
|  | 5 | 0,179 | 0,247 | 0,051 | 0,103 | 0,319 | 0,149 | 0,270 | 0,017 |
|  | Total | 0,507 | 0,629 | 0,580 | 0,530 | 0,760 | 0,495 | 0,774 | 0,942 |
| Males | 1 | 0,105 | 0,296 | 0,014 | 0,223 | 0,098 | 0,001 | 0,001 | 0,904 |
|  | 2 | 0,187 | 0,049 | 0,629 | 0,027 | 0,229 | 0,003 | 0,586 | 0,005 |
|  | 3 | 0,009 | 0,093 | 0,032 | 0,010 | 0,362 | 0,003 | 0,096 | 0,023 |
|  | 4 | 0,006 | 0,072 | 0,054 | 0,066 | 0,189 | 0,164 | 0,065 | 0,007 |
|  | 9 | 0,163 | 0,185 | 0,085 | 0,119 | 0,008 | 0,190 | 0,032 | 0,003 |
|  | Total | 0,470 | 0,693 | 0,814 | 0,446 | 0,887 | 0,361 | 0,780 | 0,943 |
| Juveniles | 1 | 0,178 | 0,086 | 0,005 | 0,062 | 0,494 | 0,027 | 0,260 | 0,922 |
|  | 2 | 0,013 | 0,307 | 0,608 | 0,158 | 0,133 | 0,184 | 0,524 | 0,018 |
|  | 3 | 0,017 | 0,269 | 0,006 | 0,079 | 0,120 | 0,271 | 0,016 | 0,002 |
|  | 7 | 0,038 | 0,043 | 0,038 | 0,103 | 0,153 | 0,079 | 0,034 | 0,000 |
|  | Total | 0,247 | 0,705 | 0,655 | 0,402 | 0,899 | 0,561 | 0,834 | 0,941 |

Highlighted cells represent cases where greater than 10% of the variation in a trait is explained by a MEM

**Table S8:** Summary of important MEM *R^2^* values of straling morphological traits adjusted by environmental conditions in Australia.

| Dataset | MEM | wingLoad | RwingL | RlegL | RbillL | RbillD | RbillW | RheadL | BR | HR | TR |
| --- | --- | --- | --- | --- | --- | --- | --- | --- | --- | --- | --- |
| All | 1 | 0,546 | 0,403 | 0,614 | 0,599 | 0,716 | 0,035 | 0,083 | 0,315 | 0,033 | 0,001 |
|  | 2 | 0,006 | 0,019 | 0,006 | 0,121 | 0,038 | 0,086 | 0,010 | 0,010 | 0,094 | 0,089 |
|  | 3 | 0,051 | 0,148 | 0,015 | 0,010 | 0,026 | 0,416 | 0,191 | 0,202 | 0,005 | 0,029 |
|  | 5 | 0,147 | 0,153 | 0,126 | 0,012 | 0,001 | 0,000 | 0,003 | 0,006 | 0,387 | 0,273 |
|  | 6 | 0,077 | 0,091 | 0,005 | 0,004 | 0,009 | 0,003 | 0,164 | 0,000 | 0,204 | 0,287 |
|  | Total | 0,828 | 0,814 | 0,766 | 0,746 | 0,790 | 0,541 | 0,450 | 0,533 | 0,724 | 0,681 |
| Females | 1 | 0,638 | 0,540 | 0,144 | 0,028 | 0,640 | 0,003 | 0,594 | 0,003 | 0,125 | 0,315 |
|  | 3 | 0,016 | 0,061 | 0,000 | 0,448 | 0,019 | 0,004 | 0,000 | 0,219 | 0,002 | 0,002 |
|  | 5 | 0,057 | 0,157 | 0,510 | 0,167 | 0,086 | 0,527 | 0,001 | 0,388 | 0,419 | 0,223 |
|  | Total | 0,711 | 0,759 | 0,654 | 0,643 | 0,745 | 0,534 | 0,595 | 0,610 | 0,546 | 0,540 |
| Males | 1 | 0,440 | 0,438 | 0,321 | 0,058 | 0,217 | 0,060 | 0,072 | 0,243 | 0,014 | 0,039 |
|  | 2 | 0,148 | 0,052 | 0,079 | 0,156 | 0,150 | 0,046 | 0,446 | 0,007 | 0,494 | 0,267 |
|  | 5 | 0,159 | 0,272 | 0,073 | 0,022 | 0,282 | 0,005 | 0,062 | 0,047 | 0,000 | 0,089 |
|  | Total | 0,747 | 0,762 | 0,473 | 0,235 | 0,649 | 0,111 | 0,580 | 0,298 | 0,509 | 0,395 |
| Juveniles | 1 | 0,005 | 0,104 | 0,541 | 0,881 | 0,658 | 0,053 | 0,229 | 0,651 | 0,009 | 0,001 |
|  | 2 | 0,001 | 0,000 | 0,093 | 0,000 | 0,008 | 0,220 | 0,267 | 0,083 | 0,048 | 0,157 |
|  | 3 | 0,002 | 0,081 | 0,016 | 0,026 | 0,008 | 0,414 | 0,012 | 0,090 | 0,000 | 0,001 |
|  | 7 | 0,148 | 0,229 | 0,035 | 0,007 | 0,015 | 0,007 | 0,074 | 0,000 | 0,350 | 0,306 |
|  | Total | 0,156 | 0,413 | 0,684 | 0,913 | 0,689 | 0,694 | 0,582 | 0,823 | 0,408 | 0,466 |
| Highlighted cells represent cases where greater than 10% of the variation in a trait is explained by a MEM |  |  |  |  |  |  |  |  |  |  |  |

**Figure S1:**
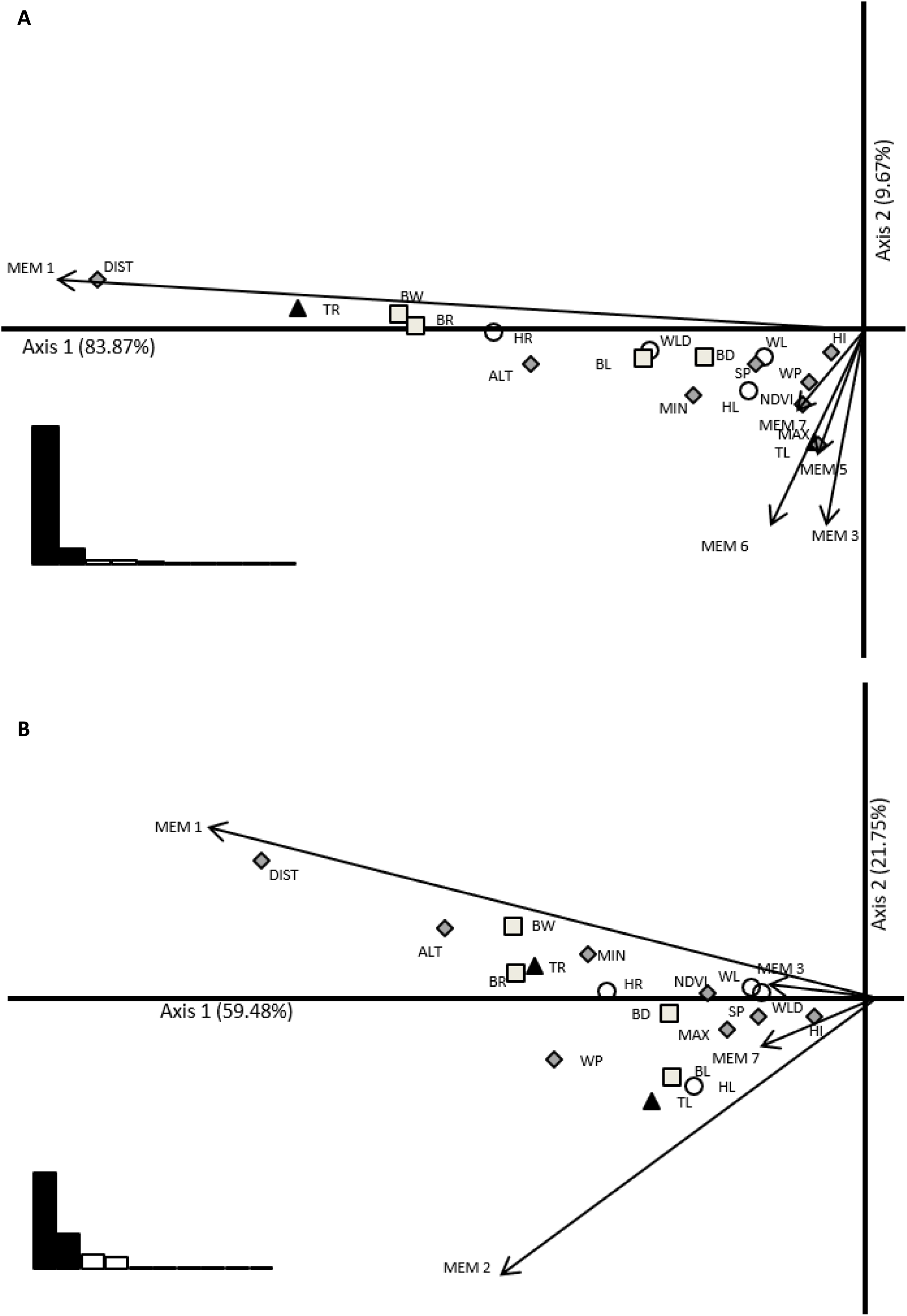
Results of environmental and morphometric analysis using MSPA redundancy analysis for the South African dataset. Eigen values are shown as insets and MEMs indicate predictors of spatial scales. A: All Data and B: Juveniles. Filled squares represent traits related to bill morphology, filled triangles traits related to tarsus, open circles traits related to dispersal, and filled diamonds environmental variables. WL: wing length, WLD: wing loading, HL: head length, HR: head ration, BW: bill width, BL: bill length, BD: bill depth, BR: bill ratio, TL: tarsus length, TR: tarsus ratio, DIST: distance from the introduction site, ALT: altitude, MIN: minimum temperature, MAX: maximum temperature, SP: mean summer precipitation, WP: mean winter precipitation, NDVI: normalised difference vegetation index AND HI: human impact.

**Figure S2:**
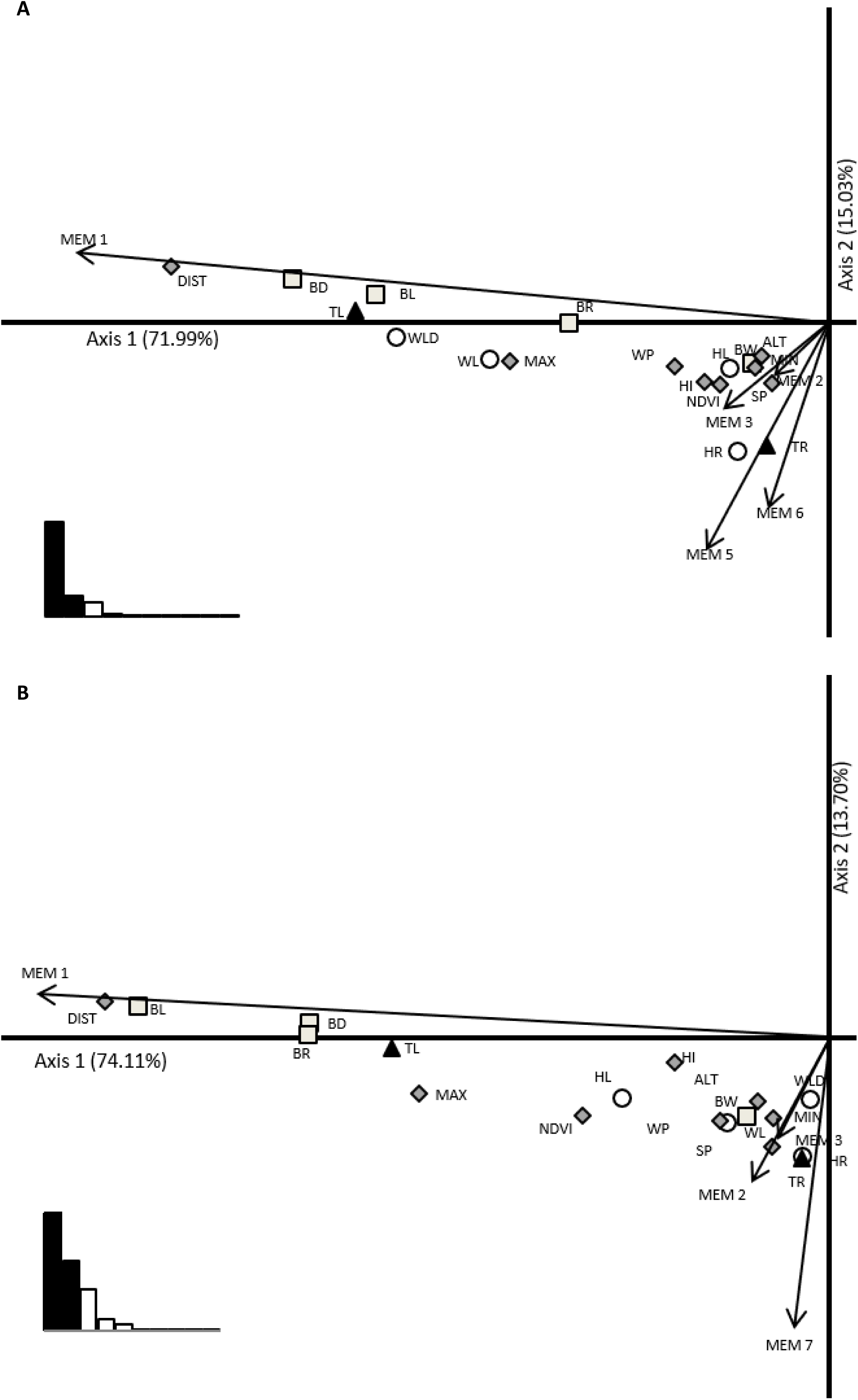
Results of environmental and morphometric analysis using MSPA redundancy analysis for the Australian Dataset. Eigen values are shown as insets, MEMs indicate predictors of spatial scales. A: All Data and B: Juveniles. Filled squares represent traits related to bill morphology, filled triangles traits related to tarsus, open circles traits related to dispersal, and filled diamonds environmental variables. WL: wing length, WLD: wing loading, HL: head length, HR: head ration, BW: bill width, BL: bill length, BD: bill depth, BR: bill ratio, TL: tarsus length, TR: tarsus ratio, DIST: distance from the introduction site, ALT: altitude, MIN: minimum temperature, MAX: maximum temperature, SP: mean summer precipitation, WP: mean winter precipitation, NDVI: normalised difference vegetation index AND HI: human impact.

